# Interdimer “zipping” in the chemoreceptor signaling domain revealed by molecular dynamics simulations

**DOI:** 10.1101/745117

**Authors:** Marharyta G. Petukh, Davi R. Ortega, Jerome Baudry, Igor B. Zhulin

**Affiliations:** Biology Department, Presbyterian College, Clinton, SC 29325; Department of Biology and Biological Engineering, California Institute of Technology, Pasadena, CA 91125; Department of Biological Sciences, The University of Alabama, Huntsville, AL 35899; Department of Microbiology, The Ohio State University, Columbus, OH 43210

## Abstract

Chemoreceptors are principal components of the bacterial sensory system that modulates cellular motility. They detect changes in the environment and transmit information to CheA histidine kinase, which ultimately controls cellular flagellar motors. The prototypical Tsr chemoreceptor in *E. coli* is a homodimer containing two principal functional modules: (i) a periplasmic ligand-binding domain and (ii) a cytoplasmic signaling domain. Chemoreceptor dimers are arranged into a trimer of dimers at the tip of the signaling domain comprising a minimal physical unit essential for enhancing the CheA activity several hundredfold. Trimers of dimers are arranged into highly ordered hexagon arrays at the cell pole; however, the mechanism underlying the trimer-of-dimer and higher order array formation remains unclear. Furthermore, molecular mechanisms of signal transduction that are likely to involve inter-dimer interactions are not fully understood. Here we apply all-atom, microsecond-time scale molecular dynamics simulations of the Tsr trimer of dimers atomic model in order to obtain further insight into potential interactions within the chemoreceptor signaling unit. We show extensive interactions between homodimers at the hairpin tip of the signaling domain, where strong hydrophobic interactions maintain binding. A subsequent zipping of homodimers is facilitated by electrostatic interactions, in particular by polar solvation energy and salt bridges that stabilize the final compact structure, which extends beyond the kinase interacting subdomain. Our study provides evidence that interdimer interactions within the chemoreceptor signaling domain are more complex than previously thought.

## INTRODUCTION

Many motile bacteria actively navigate in their environment by detecting changes in various physico-chemical parameters and controlling the direction of flagellar rotation, so they swim in a favorable direction – towards nutrients and away from toxins. Changes in the microenvironment are detected by chemoreceptors, also known as methyl-accepting chemotaxis proteins (MCPs) (1-3). MCPs detect various environmental cues (primarily small molecules) and modulate the activity of a histidine kinase CheA, which, following autophosphorylation, donates the phosphate group to a cognate response regulator CheY. In a phosphorylated form, CheY binds to a flagellar motor, which increases the probability of clockwise rotation leading to reorientation of the swimming direction (4). The serine chemoreceptor Tsr in *Escherichia coli* is one of the best studied MCPs and it serves as a model for elucidating signal transduction by various experimental and computational approaches (5-11). Similarly to other chemoreceptors, the Tsr structural unit is a homodimer, where each protomer contains a periplasmic ligand-binding domain, two transmembrane helices, a HAMP domain, and an intracellular signaling domain (Fig. 1). While ligand-binding domains in chemoreceptors are extremely diverse (12), the signaling domain is well conserved (13) – it exhibits a long hairpin supercoil structure formed by two protomers made of two antiparallel helices (14). Chemoreceptor homodimers interact at the tip of the signaling domain to form trimers of dimers. Together with CheA and an adaptor protein CheW, chemoreceptor trimers of dimers form higher order clusters of signaling complexes that are arranged as an extended lattice – a hexagonal array, which is conserved among different transmembrane chemoreceptors and various bacterial species (15-19).

**Figure 1.**
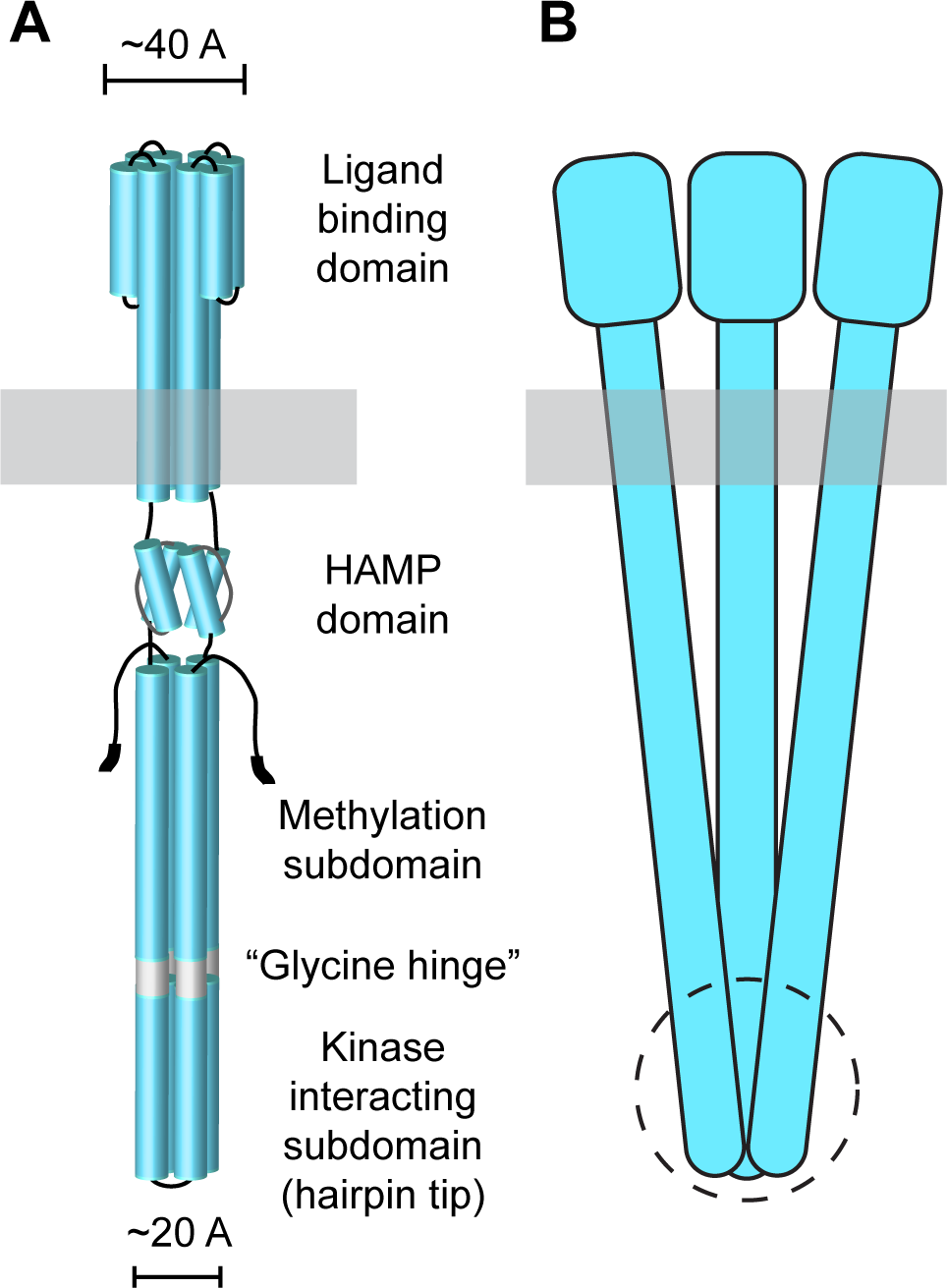
Cartoon model of the *E. coli* Tsr chemoreceptor. (A) The Tsr homodimer consists of alpha-helical domains: ligand-binding domain, transmembrane regions, HAMP domain, and signaling domain organized in two subdomains (methylation and kinase interacting) separated by the glycine hinge. The subunits of each dimer form a four-helix bundle. (B) Proposed arrangement of the Tsr trimer of dimers. All three dimers in the trimer interact at the hairpin tip.

Signaling complexes exhibit two-state kinetics: high CheA activity generates the “ON” state, whereas low CheA activity leads to the “OFF” state (1). Autophosphorylated CheA donates the phosphate group not only to CheY, but also to a CheB methylesterase (albeit with a much slower rate). Phosphorylated CheB removes methyl groups that are constitutively added by a CheR methyltransferase to several methylation sites on the MCP signaling domain. The CheB-CheR pair comprises the sensory adaptation system functioning as a negative feedback loop. The cluster organization of chemoreceptors contributes to the high sensitivity, wide dynamic range, and precise adaptation of the signaling system (20, 21). The current view of the trimer of dimers assembly is based on the crystallographic structure of the Tsr trimer of dimers (14), disulfide mapping (22), and cryoelectron tomography of chemoreceptor arrays (17, 18, 23): it is formed through extensive interactions between dimers throughout the kinase interacting subdomain, however, dimers do not interact with each other above this subdomain (Fig. 1a). Consequently, the dimers cross the membrane at a certain angle avoiding structural clashes between chemoreceptor periplasmic domains that are much wider than cytoplasmic ones (Fig. 1b).

The process of signal transduction by chemoreceptors is expected to be highly dynamic. It likely involves various types of conformational changes: regional and global motions of the chemoreceptor dimer as well as intra- and interdimer interactions are thought to be important for signal transduction (24-28). The overall chemoreceptor structure is rigid; however, two flexible regions were identified in the cytoplasmic part of the protein. The “glycine hinge” (Fig. 1a) is comprised of conserved glycine residues in each protomer and allows the four-helix bundle of the dimer to bend ∼ 10 degrees (24). Molecular dynamics simulation of the full-length Tsr dimer model revealed the second potential “hinge”, just below the HAMP domain, which potentially can allow ∼30 degrees bending of the cytoplasmic four-helix bundle (29). A recent imaging of intact, lipid-bilayer-inserted chemoreceptor dimers provided further support for two flexible hinges in the signaling domain (30). However, the prevalent view is that dynamic changes within and between signaling complexes are expected to be relatively small, because arrays of chemoreceptors in different signaling states exhibit the same architecture in electron cryotomogramms (31, 32).

To obtain further insight into potential interactions within the chemoreceptor signaling unit, we performed all-atom, microsecond-range molecular dynamics (MD) simulations of the Tsr signaling domain in its trimer of dimers state. Here, we report extensive interactions between homodimers within the trimer, describe their nature and discuss potential implications of the observed dynamic behavior.

## METHODS

### MD simulation system

The initial simulation system was built from the model of the Tsr signaling domain dimer obtained from the last frame of the previously described 2 µs production simulation (28). To build the trimer of dimers, we aligned three copies of the initial model with each dimer of the crystal structure of the Tsr chemoreceptor (PDB ID: 1QU7) (14). Structural alignment was done using STAMP from the VMD software (33) with default parameters. The model was then charge-neutralized, supplemented with 150 mM of NaCl, and embedded in a water (TIP3P) hydration box. The total simulation system contained 347,829 atoms in a 127 x 135 x 235 Å periodic box. The system was energy-minimized using the conjugate gradient algorithm for 400 steps and equilibrated in 1 ns MD simulation at T=298 K and P=1 atm using the CHARMM27 (34) force fields, NPT ensemble, Nose-Hoover Langevin piston algorithm (35) with a period of 100 fs, a decay time of 50 fs, and a 2 fs integration time step with a multiple time stepping scheme (36). All van der Waals forces were calculated with a cutoff value of 12 Å, and a switching function starting at a 10 Å distance. Electrostatic interactions were calculated using a particle mesh Ewald (PME) algorithm. Energy minimization and the molecular dynamics simulation (MD) equilibration steps were performed on the Newton computing cluster at The University of Tennessee-Knoxville. A one µs production MD simulation was performed on the Titan supercomputer in Oak Ridge National Laboratory using NAMD2 v2.9 (37)

### Residues forming interface between interacting dimers

To quantify inter-dimer interactions, we calculated the difference in the relative solvent accessible surface area (SASA) of the residues in the single dimer unit and in the dimer-of-dimers observed during the simulation:

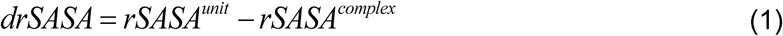

In eq. (1) *rSASA^unit^* is the relative SASA of the residue being in a single (AB), (CD), or (EF) dimer, while *rSASA^complex^* is the relative SASA of the residue when the complex of two dimers (i.e. (AB)-(CD), (AB)-(EF), or (CD)-(EF)) was formed. Thus, the non-zero value of *drSASA* of a residue indicates that it is at the dimer interface, participating in interdimer interactions.

Residues engaging in hydrogen bond interactions between their side-chains were identified based on the following criteria: the distance between the hydrogen donor and its acceptor is within 3.0±0.4 Å and acceptor-hydrogen-donor angle is in the 180°±20°range. Oxygen, nitrogen, and sulfur atoms were considered for both acceptor and donor of hydrogen atom. Cases where the acceptor-hydrogen-donor pairs can establish a salt bridge were not included in a hydrogen bond count. The VMD software (33) was used to identify residue pairs forming salt bridges. Standard parameters for salt bridge geometry were used with a distance cutoff between nitrogen atoms of positively charged residues and oxygen atoms of the COO^−^ moiety in negatively charged residues set at 4 Å. Hydrophobic interactions between Ala, Val, Leu, Ile, Phe, Trp, and Met residues were identified in the MD trajectory by calculating the distance between sidechain atoms. The cutoff distance between any pairs of sidechain atoms was set at 4 Å.

### Energetic characteristics of dimer-of-dimers zipping

To describe the system behavior during MD simulations, we performed energy calculations. The energy deviation of the dimer-of-dimers system 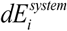 from its average value at the initiation state (eq.(2)) was calculated every 0.2 ns during the MD simulations. The data were normalized by the absolute value of the average energy at the initiation state.

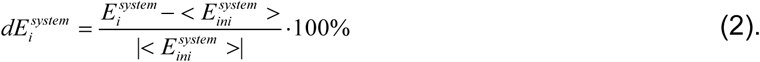

The second type of calculations was performed at the equilibrium state only. It reveals the preference of the dimer-of-dimers system to stay in the zipped state compared to a completely unbound state 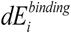 (eq.(3)) In eq. (3) the negative value of 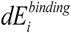 points out that the system favors the bound state over the unbound; the opposite is valid for a positive value of the 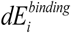. The magnitude indicates the degree at which one of the states – bound or unbound – is more preferable than another.

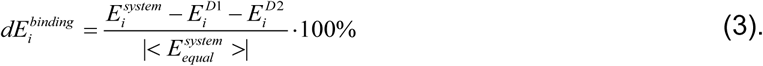

In eq. (3) 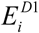 and 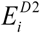 are the energies of the (CE) and (EF) dimers within the complex. For simplicity, these energies were calculated for the dimers being structurally identical to the one in the complex without taking into consideration potential structural changes due to interdimer interactions.

We considered the following energy contributions: mechanic (ME), van der Waals (VE), non-polar solvation (SN), coulombic (EE), and polar solvation (SP). ME, EE and VE energy terms were obtained during one step minimization with NAMD (37). ME was determined by the sum of bond, angle, dihedral, and improper components of energy.

The SN was estimated via linear formula with respect to SASA of the system:

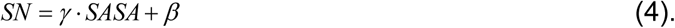

In eq. (4) *γ* = 0.00542 kcal/mol/Å^2^ and *β* = 0.92 kcal/mol (38). The SASA was calculated with NACCESS software (39).

The SP energy component was calculated with Delhi software (40) with the standard parameters: scale 1 Å per grid, perfil 70 %, dielectric constants for protein was 1 and water – 80.

### Hydrophobicity and entropy

The entropy of the system was estimated with a previously described empirical formula (41, 42), based on the maximal number of side chain rotamers (R) (43):

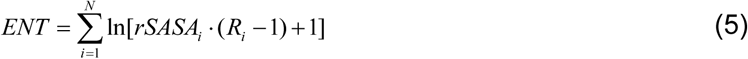

The hydrophobicity of the system of N residues was estimated with a previously described empirical formula (41, 42) using relative solvent accessibility (*rSASA*) of each residue and its hydrophobicity characteristics (*H*) assigned by Wimley-White hydrophobicity scale (44):

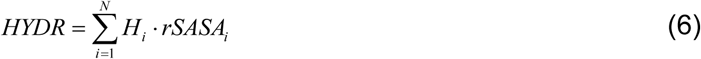

### Sequence conservation

Conservation of each amino acid position within the Tsr signaling domain was assessed by constructing a multiple sequence alignment of the signaling domain regions from 535 homologous chemoreceptors from enterobacterial genomes, as previously described (45). The MiST database (46) was the primary source of sequence data including pre-computed features (e.g. domains, transmembrane regions, etc.). The following two criteria were applied in selecting the chemoreceptor sequences for analysis: (i) sequences must belong to Type I membrane topology class, i.e. contain a periplasmic ligand-binding domain between two transmembrane helices (47) and (ii) sequences must belong to the 36H length class, i.e. contain 36 helical heptads in the signaling domain (13). A multiple sequence alignment was constructed using L-INSI-I algorithm from MAFFT (48) and amino acid conservation was calculated using the CONSENSUS script available at http://consensus.zhulinlab.org/. Conservation of residues within the kinase interacting subdomain was assessed from a previously published dataset of nearly 8000 MCP sequences from diverse bacterial and archaeal phyla (28).

## RESULTS

### Interdimer Zipping

The structure of the Tsr signaling domain (PDB ID 1QU7) trimer comprises three homodimers – (AB), (CD), and (EF) – connected only through the tips of their signaling domains (Fig. 2a). After ∼400 ns of MD simulations, two of the dimers, (CD) and (EF), bound together throughout their full length, whereas the (AB) dimer stood apart, interacting with the other two dimers only through the tip (Fig. 2). The timescale of the oligomerization was similar to that previously observed in a coarse-grain MD simulation of the chemoreceptor trimer of dimers model (29).

**Figure 2.**
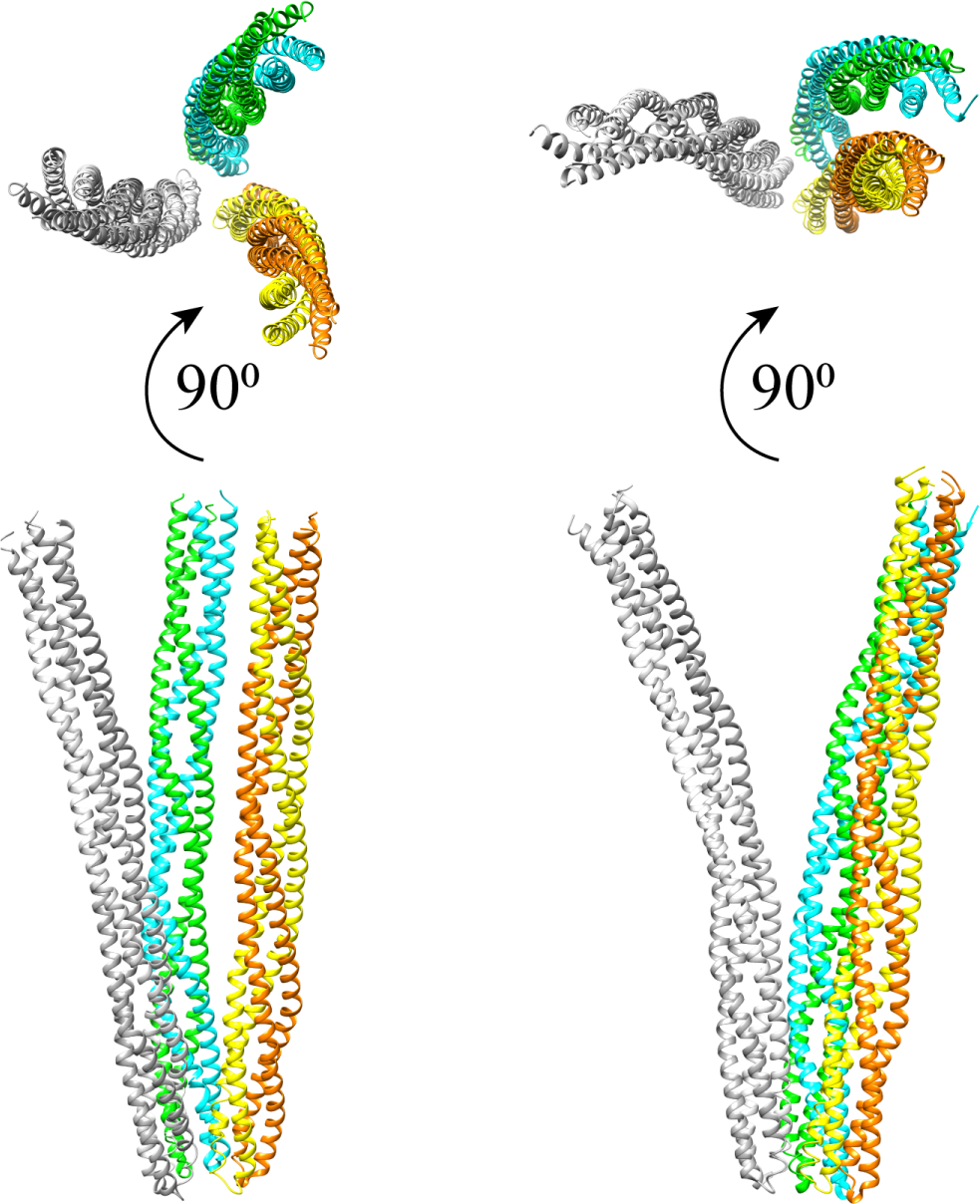
Two-dimer interaction within the signaling domain trimer of dimers. Initial (left) and final (right) snapshots of the Tsr signaling domain trimer of dimers one-microsecond MD simulation are shown as top and side views. Monomers comprising (AB) dimer are shown in light and dark grey, monomers comprising (CD) dimer are shown in yellow and orange, and monomers comprising (EF) dimer are shown in cyan and green.

The RMSD analysis suggested three phases of the interdimer interaction: (i) initiation, the first 100 ns; (ii) “zipping”, the next ∼300 ns, which is defined by a significant increase in the RMSD value; and (iii) equilibrium, from 400 ns until the end of MD simulations (Fig. 3A). The nearly constant RMSD of zipped (CD) and (EF) dimers at the equilibrium phase indicates that the motion of each dimer is significantly reduced. Substantial fluctuations in the RMSD of the full system (Fig. 3) are likely due to the lack of constraints on the unzipped (AB) dimer. The amplitude of the residue motions within all six protomers at the equilibrium state was similar (Fig. S1). The residues fluctuate the most at both N- and C-termini, as well as at the chemoreceptor tip, similarly to the behavior previously described for an isolated Tsr dimer (49). To quantify the dynamics of (CD)-(EF) zipping, we monitored the total number of contacting residues that constitute the zipped interface. The average number of interface residues increased from 33 at the initiation phase to 113 at the equilibrium phase, with a zipping speed of ∼0.6 residue per ns (Fig. 3B).

**Figure 3.**
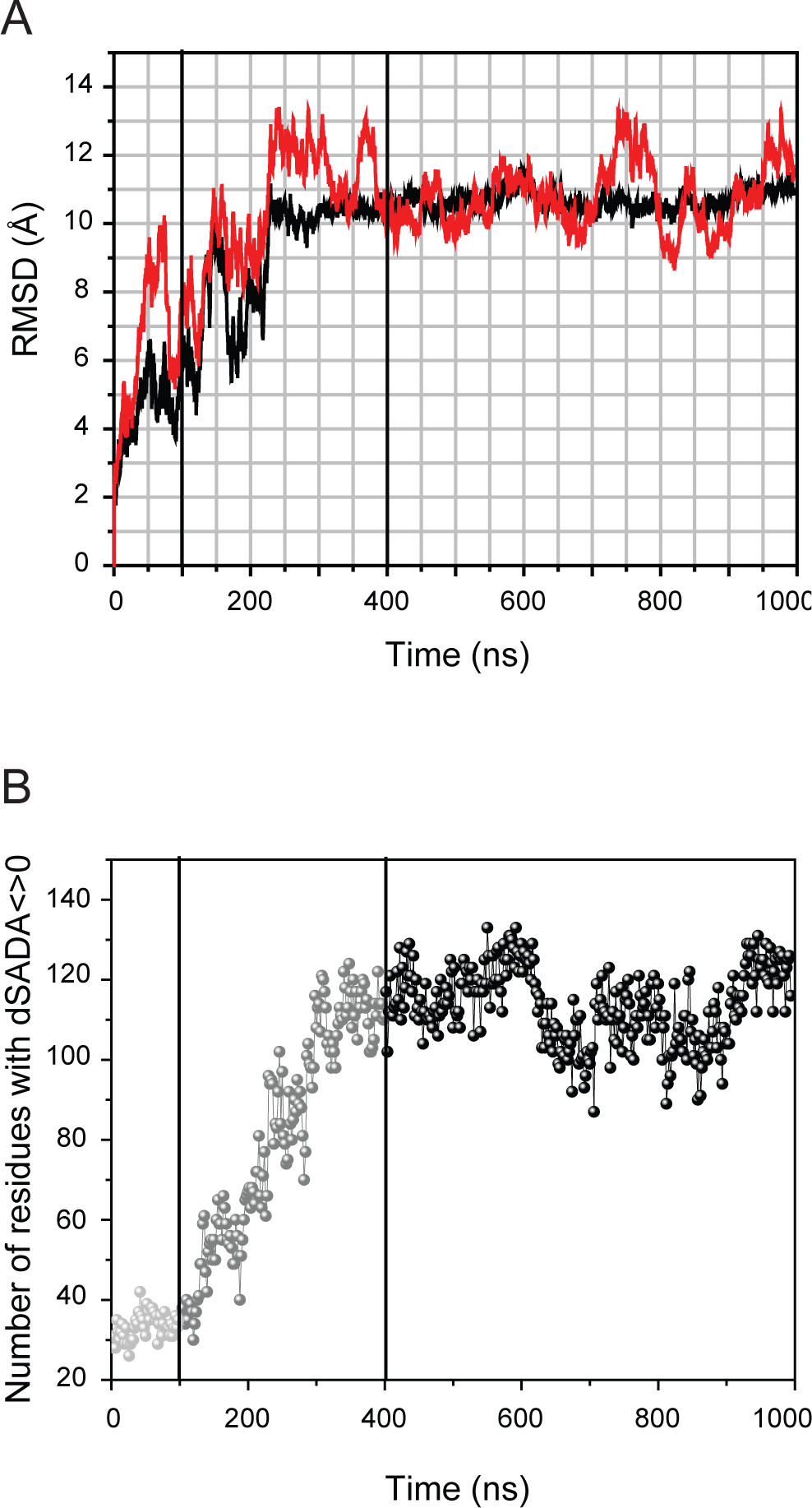
Dynamics of the Tsr signaling domain inter-dimer interactions. (A) RMSD values for the (CD)-(EF) dimer (black line) and the (AB)-(CD)-(EF) trimer (red) monitored along 1 µs MD simulation. (B) Number of contacting residues between (CD) and (EF) dimers during MD trajectory.

To characterize how the residues of (CD) and (EF) homodimers interact during the zipping phase, we calculated the change in the *rSASA* of the residues within each plane (a cross-section of each homodimer consisting of two identical pairs of residues from two protomers) of both (CD) and (EF) homodimers during the simulation. We found that while zipping propagated from the chemoreceptor tip (a plane containing Gln 392 at the hairpin turn) to the N and C termini of the model, it occurred in a non-sequential residue order. The system sampled multiple interaction sites by establishing temporal interdimer interactions, breaking some (presumably redundant) and preserving energetically favorable ones (Fig. S2). The final binding sites were established at the equilibrium stage, as the *drSASA* of residues in protomers C, D, E, and F did not dramatically change during the last 600 ns of MD simulation. Interacting residues on each protomer were distributed in two to three patches throughout the entire length of the signaling domain (Fig. S3). Notably, residues that comprise methylation sites were not involved in interdimer interactions and remained solvent-exposed throughout the MD simulation (Fig. S3).

### Zipping is driven by electrostatics

We calculated values of several energy components and the total hydrophobicity parameter of the (CD)-(EF) interaction system during the MD simulation and observed significant reduction in polar/nonpolar solvation energy components and in the hydrophobicity parameter, specifically during the zipping phase (Fig. S4). Analysis of the binding free energy in relation to the total energy of the (CD)-(EF) complex shows that the system released ∼20% of polar solvation component and ∼10% of nonpolar solvation component of the total energy due to interdimer binding (Fig. S5). The decrease in the hydrophobicity parameter indicates that during zipping the number of solvent-exposed hydrophilic residues was reduced, likely due to establishing interdimer salt bridges and hydrogen bonds. Considering that each of the homodimers has a large absolute total charge of 26, the unzipped pair of dimers incurs a significant desolvation penalty. The change in solvation due to binding can be explained by the Born effect that is observed to guide binding of highly charged protomers with the same charge polarity (50).

We determined how specific bonds – hydrophobic interactions, salt bridges, and hydrogen bonds – between homodimers changed over the time and calculated the frequency of specific bonds formation at the equilibrium phase. At the initiation phase, interdimer interactions occurred only at the chemoreceptor tip, where, on average, seven hydrophobic contacts between (CE) and (DF) dimers were detected. During the zipping phase, only one or two additional hydrophobic contacts were gained and this remained unchanged throughout the equilibrium phase (Fig. 4a). Stability of hydrophobic interactions was further confirmed by high “frequency of participation” - the amount of time a given residue spends in a specific interaction with another residue (Table S1). There were only two salt bridges (on average) in the (CE)-(DF) complex at the initiation phase, but their number during the zipping phase linearly increased reaching the average of eight at the equilibrium phase (Figure 4b and Fig. S6) and showing high frequency of participation (Table S2). Only a single hydrogen bond between homodimers was detected during the initiation phase (Fig. S7). During the zipping phase, the number of hydrogen bonds increased to up to six, but it fluctuated extensively in the equilibrium stage. The statistical analysis shows that the average number of hydrogen bonds at the equilibrium stage was two, and the large standard deviations of the average suggest that the hydrogen bonds are unstable. This is further confirmed by the low extent of residue participation in interdimer hydrogen bond formation (Table S3).

**Figure 4.**
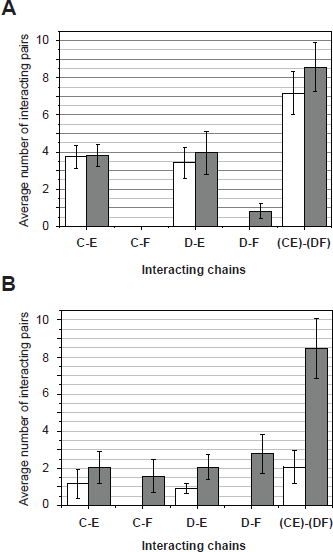
Comparison of intra- and inter-dimer contacts at initiation and equilibrium stages. (A) Hydrophobic interactions. (B) Salt bridges. White, initiation; grey, equilibrium.

### Interdimer contacts at the chemoreceptor tip

Previously, several interdimer contacts had been deduced from the crystal structure of the Tsr trimer of dimers, all within the chemoreceptor tip (14). Specifically, three hydrophobic contacts (Phe 373 – Ile 377, Leu 380 – Leu 378, and Val 384 – Val 398), one salt bridge (Glu 385 – Arg 388) and two hydrogen bonds (Asn 376 – Arg 409 and Asn 381 – Asn 381’) were proposed. More recently, another salt bridge (Glu 402 – Arg 404) was identified and shown to stabilize the Tsr trimer of dimers (10). All participating residues were previously identified as highly conserved. Our data largely agree with these results, but also show that interdimer interactions within the chemoreceptor tip are more complex than previously thought. First, we found that some hydrophobic residues make contacts with more than one residue: twelve contact pairs are formed by only twelve residues (Table 1). Ile 377, Leu 380, and Val 398, each participate in three contact pairs. Six contact pairs are formed between interacting N (C) – N (E) helices, five pairs are between C (D) and N (E) helices, and a single pair (Glu 402 – Arg 404 salt bridge) is between C (D) and C (E) helices (Fig. 5). This intertwined web of contacts and high incidence of interacting hydrophobic residues indicate very strong interdimer interactions at the tip and a nearly perfect conservation of the key contact residues over very long evolutionary time is consistent with their importance.

**Table 1.** Key interdimer contacts in the kinase interacting subdomain.

| Bond |  | Type | Frequency,<br>% | Conservation,<br>% |  |
| --- | --- | --- | --- | --- | --- |
| Residue 1 | Residue 2 |  |  | Residue 1 | Residue 2 |
| F373 (C) | F373 (E) | Hydrophobic | 99 | 33 | 33 |
| I377 (C) | F373 (E) | Hydrophobic | 96 | 100 | 33 |
| I377 (C) | I377 (E) | Hydrophobic | 64 | 100 | 100 |
| L378 (C) | I377 (E) | Hydrophobic | 27 | 100 | 100 |
| L378 (C) | L380 (E) | Hydrophobic | 87 | 100 | 100 |
| E385 (C) | R388 (E) | Salt bridge | 50 | 99 | 98 |
| V398 (D) | L380 (E) | Hydrophobic | 89 | 98 | 100 |
| V398 (D) | A383 (E) | Hydrophobic | 23 | 98 | 72 |
| V398 (D) | V384 (E) | Hydrophobic | 94 | 98 | 100 |
| V399 (D) | L380 (E) | Hydrophobic | 55 | 100 | 100 |
| V399 (D) | V384 (E) | Hydrophobic | 74 | 100 | 100 |
| E402 (D) | R404 (E) | Salt bridge | 80 | 99 | 97 |

Contacts with frequencies >20% are shown. (C), (D), and (E) specify the protomer, where a given residue is located. Frequency shows % of time residue 1 interacts with residue 2. Conservation of a chemical property necessary to maintain a contact (hydrophobicity, relevant charge) is derived from a multiple sequence alignment of nearly 8,000 chemoreceptor sequences from diverse bacteria phyla (Dataset 1).

**Figure 5.**
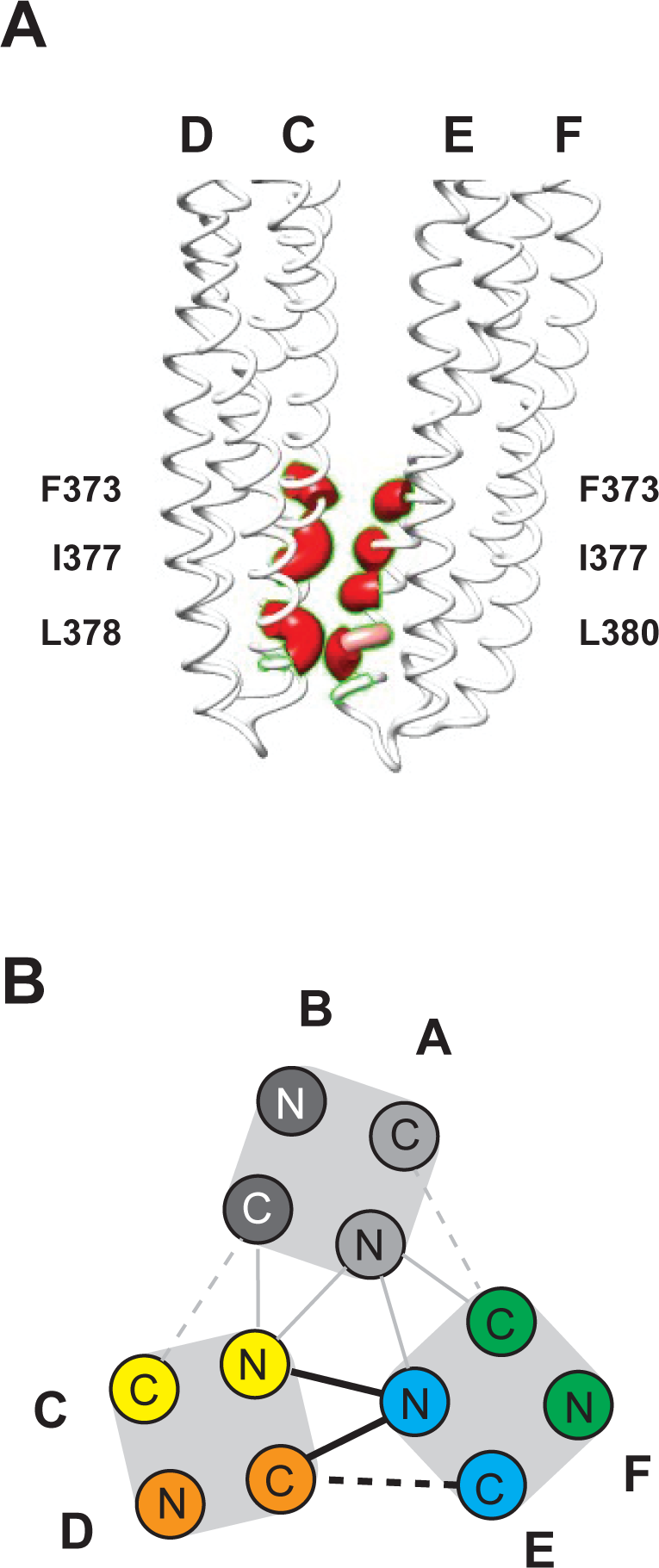
Key hydrophobic interactions in the kinase activating subdomain. (A) Residues participating in several pairs are shown on the (CD)/(EF) interface. (B) A plane of the trimer of dimers showing key inter-dimer interactions. Hydrophobic interactions are shown as solid lines and salt bridges as dashed lines. Measured and projected interactions are shown in black and grey lines, respectively.

### Interdimer contacts above the kinase-interacting subdomain

No interdimer contacts above the kinase-interacting subdomain were previously reported. At the equilibrium phase, thirteen contacts with frequencies >20% were identified between zipped (CD) and (EF) dimers above the kinase-interacting subdomain (Table 2). Only two contacts were via hydrophobic interactions (Ala 314 (D) – Leu 475 (F) and Met 264 (E) – Val 514 (D); those residues are poorly conserved. Eleven contacts were salt bridges. None of the residues participated in more than two contacts and only four residues (K324, K342, R482, and E510) participated in two contact pairs. In striking contrast to the pattern observed at the chemoreceptor tip, only one contact pair outside the tip (D363 - R415) had a high frequency of participation and was highly conserved at both positions. However, all contact pair residues were highly conserved in polar character suggesting high possibility of contacts in the form of weaker hydrogen bonds in cases where salt bridges cannot be formed.

**Table 2.** Key interdimer contacts outside the kinase interacting subdomain.

| Bond |  | Type | Frequency,<br>% | Conservation,<br>% |  |
| --- | --- | --- | --- | --- | --- |
| Residue 1 | Residue 2 |  |  | Residue 1 | Residue 2 |
| D269 (C) | R265 (E) | Salt bridge | 90 | 21 | 22 |
| E303 (D) | R482 (F) | Salt bridge | 68 | 64 | 23 |
| A314 (D) | L475 (F) | Hydrophobic | 69 | 62 | 43 |
| K317 (D) | D471 (F) | Salt bridge | 26 | 82 | 37 |
| R324 (D) | E321 (F) | Salt bridge | 67 | 67 | 83 |
| D363 (C) | R415 (E) | Salt bridge | 62 | 85 | 90 |
| E436 (C) | K342 (F) | Salt bridge | 20 | 37 | 23 |
| E440 (C) | K342 (F) | Salt bridge | 63 | 42 | 23 |
| E458 (C) | R324 (F) | Salt bridge | 34 | 91 | 67 |
| D481 (D) | R482 (F) | Salt bridge | 96 | 93 | 24 |
| E510 (D) | R271 (E) | Salt bridge | 62 | 26 | 83 |
| E510 (D) | R516 (F) | Salt bridge | 49 | 26 | 71 |
| V514 (D) | M264 (E) | Hydrophobic | 31 | 67 | 34 |

Contacts with frequencies >20% are shown. (C), (D), (E), and (F) specify the protomer, where a given residue is located. Frequency shows % of time residue 1 interacts with residue 2. Conservation of a chemical property necessary to maintain a contact (relevant charge, hydrophobicity) is derived from a multiple sequence alignment of enterobacterial chemoreceptors conforming to Class I membrane topology (Dataset 2).

## DISCUSSION

Despite numerous studies of the structure/function relationships within and between bacterial chemoreceptors, molecular mechanisms of signal transduction - from ligand binding to kinase activation – are not fully understood. This deficit can be attributed to the dynamic nature of signal transduction events, whereas most of the current experimental techniques provide static snapshots of various system states. Here, our data suggest that, within the cytoplasmic signaling domain, chemoreceptor dimers might interact more extensively and over longer distances than previously thought. Potentially, such extensive conformational changes can contribute to signal transduction. We have observed a phenomenon that we termed “zipping”, in which two of the three dimers within a trimer come into a close association throughout their length, with the third dimer engaging the other two dimers only at the chemoreceptor tip. The process of zipping was relatively slow (∼300 ns) and highly dynamic. Starting with strong, multiple hydrophobic contacts at the tip, zipping was governed by electrostatic interactions. First, unstable hydrogen bonds formed between the two dimers during the zipping stage, allowing the system to explore different binding modes. Then, energetically favorable conformations further stabilized through stronger and more permanent salt bridges at the equilibrium stage.

A recent cryo-ET/MD study exploring conformational changes of the Tsr signaling domain in different signaling states revealed inter-dimer interactions similar to those described here and concluded that asymmetric flexibilities within the trimer are a function of signaling state (51). However, such interactions were detected only within the kinase-interacting subdomain. We also observed strong evolutionarily conserved inter-dimer interactions within the kinase-interacting subdomain, however, in addition, our study revealed weaker inter-dimer interactions above the kinase-interacting subdomain. Inter-dimer interactions above the kinase-interacting subdomain might be prohibited in full-length receptors by physical constraints imposed by the size of the periplasmic ligand-binding domain, but this remains to be demonstrated. Moreover, recent studies have revealed other features of full-length chemoreceptors that might overcome such structural clashes: bending at two locations within the signaling domain (29, 30) and transitions between oligomeric states in the ligand-binding domain (52). Therefore, transient inter-dimer interactions above the kinase-interacting subdomain might play a signaling role in receptor core complexes.

## CONCLUSIONS

Taken together, the results of MD simulation combined with the sequence conservation analysis show that interdimer interactions within the chemoreceptor signaling domain can occur not only within, but also above the kinase-interacting subdomain. These transient and highly dynamic interactions are driven by electrostatics, which is in contrast to more permanent hydrophobic interactions that maintain a trimer of dimers at the kinase-interacting subdomain.

## AUTHOR CONTRIBUTIONS

M.G.P. conceived the study, performed computations, analyzed and interpreted data, and wrote the manuscript. D.R.O. conceived the study, performed computations, helped data analysis and interpretation, and helped write the manuscript. J.B. advised data analysis and interpretation, and helped write the manuscript. I.B.Z. conceived the study, analyzed and interpreted data, and wrote the manuscript.

## ACKNOWLEDGMENTS

We thank Keith Cassidy for discussions during the early stages of this study and Sandy Parkinson for discussions and critical reading of the manuscript.

This work was supported in part by the National Institutes of Health grant R35GM131760 to I.B.Z. This research used resources of the Oak Ridge Leadership Computing Facility at the Oak Ridge National Laboratory, which is supported by the Office of Science of the U.S. Department of Energy under Contract No. DE-AC05-00OR22725.

