## Supplementary Figures and Tables for "Interdimer “zipping” in the chemoreceptor signaling domain revealed by molecular dynamics simulations"

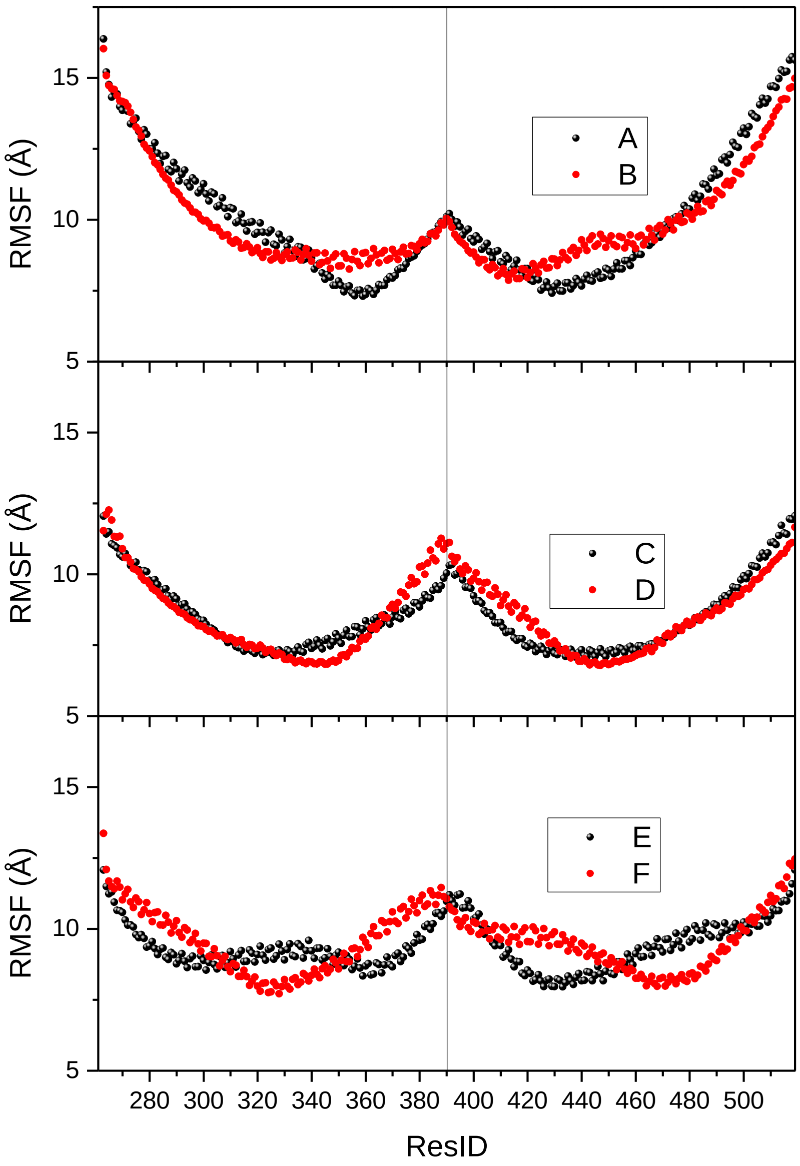


FIG. S1: **RMSF in dimers at the equilibrium stage**.

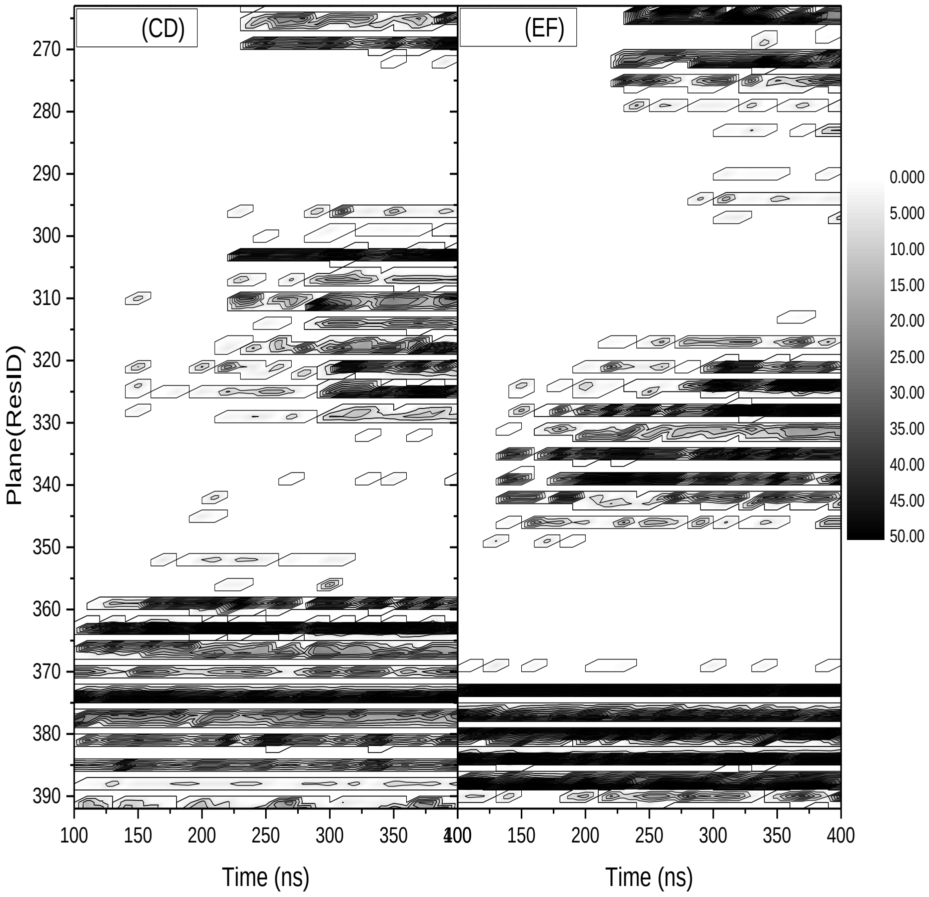


FIG S2: **Distribution of residues at the interface of dimer-of-dimers**. Left: (CD) dimer, right: (EF) dimer. On the OX-axis is the time of MD simulations. Y axis –residue number constructing the plane across protein starting from the tip of the signaling domain (plane involver ResID 392) to its terminus (plane involved ResID 263). The shades represent rSASA values of the residue due to homodimers binding.

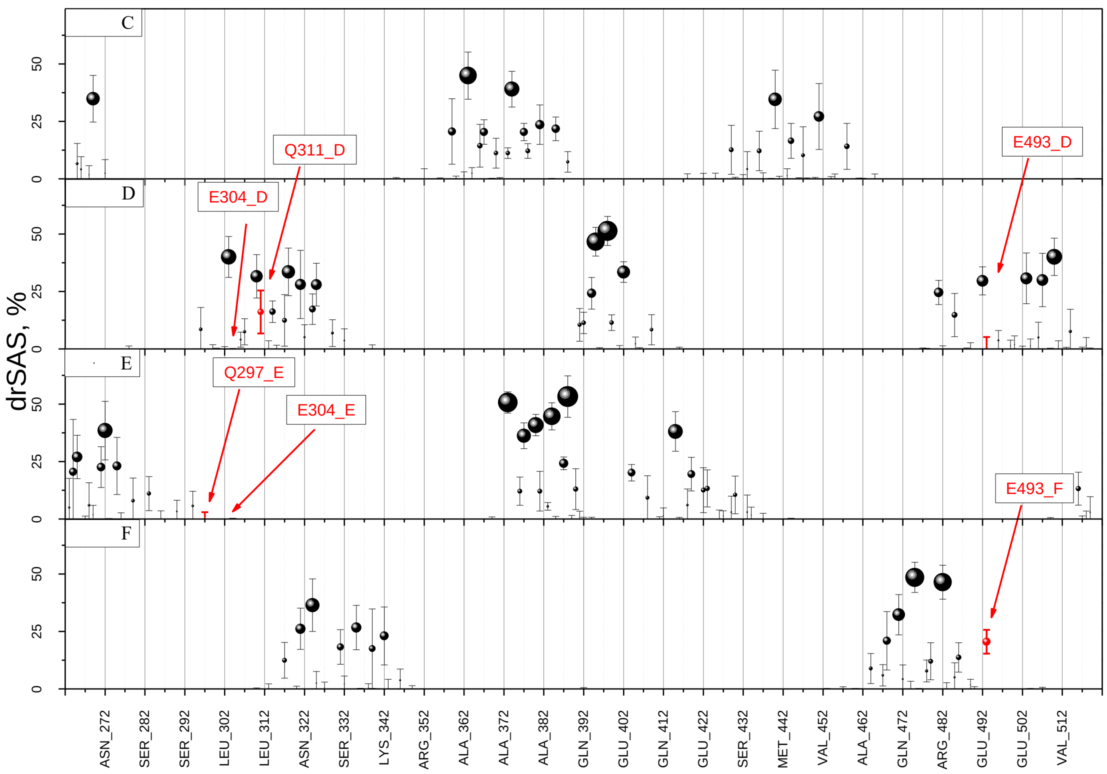


FIG S3: **Residues involved in interaction between (CD) and (EF) dimers.** X axis: residue number; Y axi:s The size of a circle is proportional to the change in the rSASA of the residue due to dimers binding. Methylation sites are indicated in red.

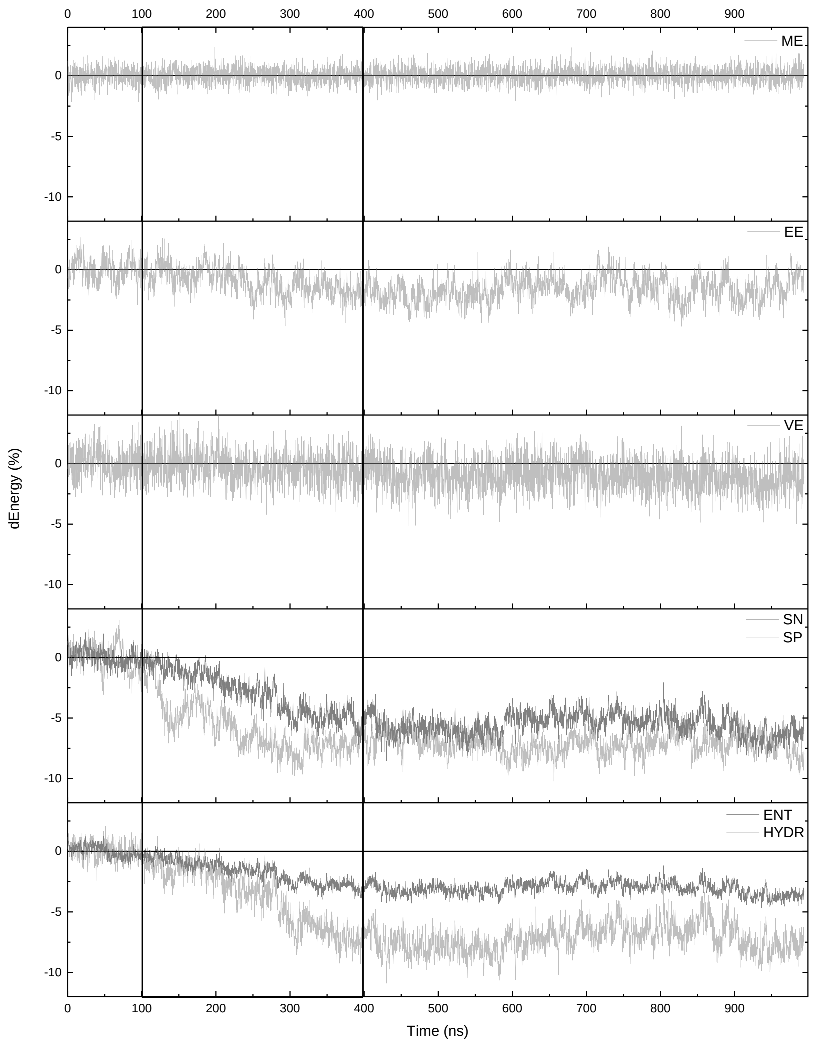


FIG S4: **The change in the energetic characteristics of the (CD)-(EF) DOD complex during MD simulation.** The data is presented for various energy components normalized by the average value of the correspondent energy term at the initiation state.

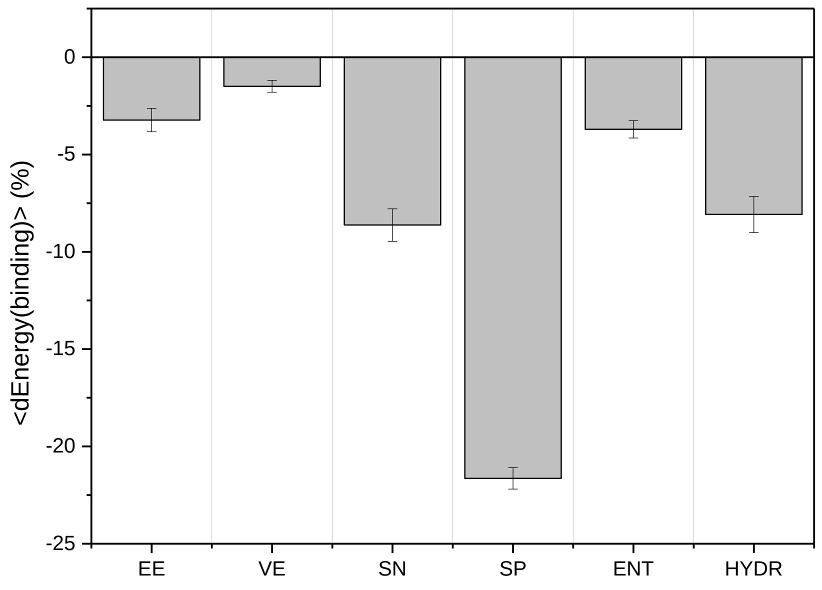


FIG S5: **The impact of different energy terms on the binding affinity between (CD) and (EF) homodimers to form a fully zipped dimer-of-dimers complex.**

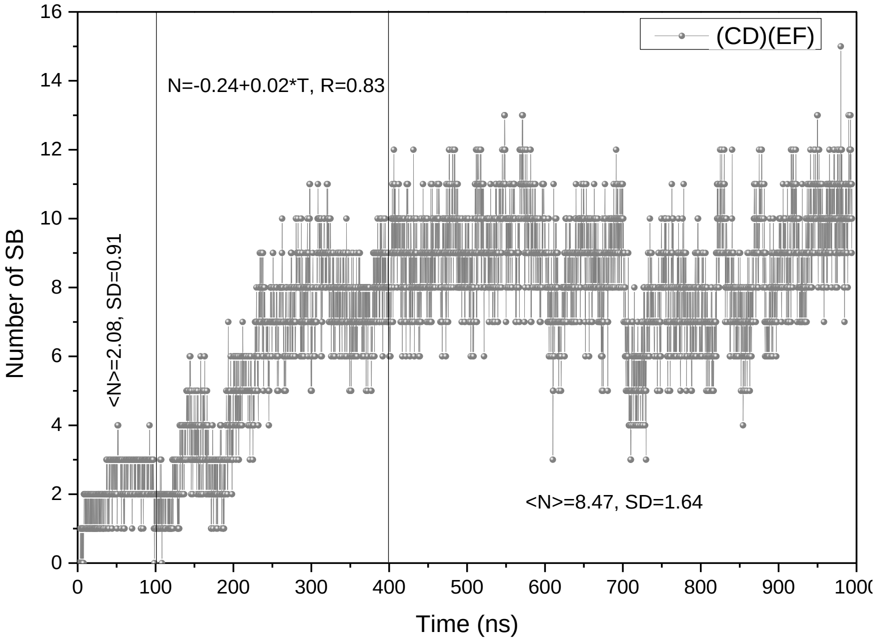


FIG S6: **Dynamics of salt bridge formation between (CD) and (EF) dimers during MD simulation.**

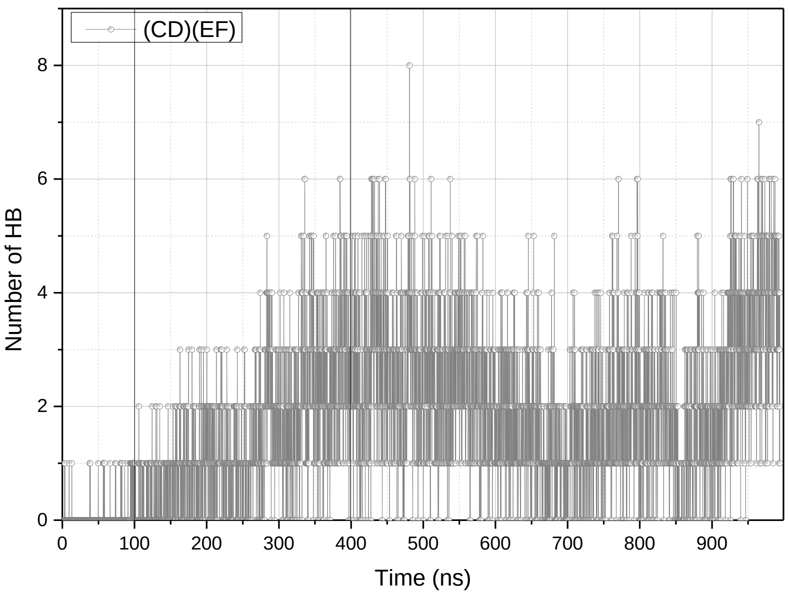


FIG S7: **Dynamics of hydrogen bond formation between (CD) and (EF) dimers during MD simulation.**

Table S1: The frequency of the residues participating in inter-dimeric hydrophobic interactions between particular monomers and the residue conservation rate based on identity and similarity (ACFGILMVW residues).

| Res | ID | Frequency, % | | | | | | | | Conservation, % | |
|---|---|---|---|---|---|---|---|---|---|---|---|
| | | C->E | C->F | D->E | D->F | E->C | E->D | F->C | F->D | Identity | Similarity |
| L | 263 | 0 | 0 | 0 | 0 | 0 | 7.98 | 0 | 0 | 89.72 | 99.81 |
| M | 264 | 0 | 0 | 0 | 0 | 2.46 | 51.18 | 0 | 0 | 1.31 | 45.79 |
| V | 270 | 2.46 | 0 | 0 | 0 | 0 | 0 | 0 | 0 | 69.91 | 100 |
| A | 276 | 0 | 0 | 0 | 0 | 0 | 0.54 | 0 | 0 | 21.12 | 26.92 |
| A | 307 | 0 | 0 | 0 | 17.36 | 0 | 0 | 0 | 0 | 89.53 | 89.72 |
| A | 314 | 0 | 0 | 0 | 63.93 | 0 | 0 | 0 | 0 | 59.07 | 63.74 |
| I | 371 | 0.27 | 0 | 0 | 0 | 0 | 0 | 0 | 0 | 96.07 | 100 |
| F | 373 | 98.86 | 0 | 0 | 0 | 197.96 | 0 | 0 | 0 | 98.88 | 99.44 |
| I | 377 | 162.96 | 0 | 0 | 0 | 91.56 | 0 | 0 | 0 | 96.26 | 100 |
| L | 378 | 114.71 | 0 | 0 | 0 | 0 | 0 | 0 | 0 | 100 | 100 |
| L | 380 | 0 | 0 | 0 | 0 | 87.28 | 147.21 | 0 | 0 | 99.63 | 100 |
| A | 383 | 0 | 0 | 0 | 0 | 0 | 23.18 | 0 | 0 | 96.26 | 96.64 |
| V | 384 | 0 | 0 | 0 | 0 | 0 | 171.94 | 0 | 0 | 96.26 | 100 |
| A | 387 | 0 | 0 | 0 | 0 | 0 | 4.55 | 0 | 0 | 99.63 | 99.63 |
| F | 396 | 0 | 0 | 3.47 | 0 | 0 | 0 | 0 | 0 | 100 | 100 |
| A | 397 | 0 | 0 | 1.35 | 0 | 0 | 0 | 0 | 0 | 93.83 | 94.21 |
| V | 398 | 0 | 0 | 209.55 | 0 | 0 | 0 | 0 | 0 | 99.44 | 100 |
| V | 399 | 0 | 0 | 129.68 | 0 | 0 | 0 | 0 | 0 | 100 | 100 |
| L | 406 | 0 | 0 | 2.83 | 0 | 0 | 0 | 0 | 0 | 100 | 100 |
| V | 430 | 0.03 | 0 | 0 | 0 | 0.03 | 0 | 0 | 0 | 4.11 | 20.75 |
| L | 475 | 0 | 0 | 0 | 0 | 0 | 0 | 0 | 81.29 | 20.93 | 45.05 |
| A | 500 | 0 | 0 | 0.54 | 0 | 0 | 0 | 0 | 0 | 27.1 | 32.52 |
| A | 511 | 0 | 0 | 18.71 | 0 | 0 | 0 | 0 | 0 | 36.33 | 66.67 |
| A | 513 | 0 | 0 | 0.07 | 0 | 0 | 0 | 0 | 0 | 33.15 | 41.95 |
| V | 514 | 0 | 0 | 38.7 | 0 | 0 | 0 | 0 | 0 | 38.01 | 67.23 |
| F | 515 | 0 | 0 | 1.75 | 0 | 0 | 0 | 0 | 0 | 99.44 | 99.81 |
| I | 517 | 0 | 0 | 0 | 0 | 0 | 0.07 | 0 | 0 | 19.1 | 94.57 |

Table S2. The frequency of the residues participating in inter-dimeric salt bridge formation between particular monomers and the residue conservation rate based on identity and similarity (any charged residue).

| Res | ID | Frequency, % | | | | | | | | Conservation, % | |
|---|---|---|---|---|---|---|---|---|---|---|---|
| | | C->E | C->F | D->E | D->F | E->C | E->D | F->C | F->D | Identity | Similarity |
| R | 265 | 0 | 0 | 0 | 0 | 89.91 | 0 | 0 | 0 | 10.47 | 42.06 |
| D | 269 | 89.91 | 0 | 0 | 0 | 0 | 1.82 | 0 | 0 | 11.21 | 28.22 |
| R | 271 | 0 | 0 | 0 | 0 | 0 | 66.49 | 0 | 0 | 72.9 | 85.42 |
| R | 294 | 0 | 0 | 0 | 0 | 0 | 11.24 | 0 | 0 | 92.9 | 93.46 |
| E | 296 | 0 | 0 | 4.41 | 0 | 0 | 0 | 0 | 0 | 87.66 | 91.78 |
| E | 303 | 0 | 0 | 6.83 | 68.47 | 0 | 0 | 0 | 0 | 62.99 | 66.73 |
| K | 317 | 0 | 0 | 0 | 26.04 | 0 | 0 | 0 | 10.36 | 66.36 | 86.73 |
| E | 321 | 0 | 0 | 0 | 22.37 | 0 | 0 | 0 | 67.33 | 31.21 | 84.11 |
| R | 324 | 0 | 0 | 0 | 67.33 | 0 | 0 | 33.51 | 12.01 | 44.11 | 74.39 |
| E | 335 | 0 | 0 | 0 | 0 | 0 | 0 | 14.6 | 0 | 19.44 | 47.66 |
| R | 339 | 0 | 0 | 0 | 0 | 0 | 0 | 16.69 | 0 | 20.93 | 61.68 |
| K | 342 | 0 | 0 | 0 | 0 | 0 | 0 | 83.28 | 0 | 18.88 | 64.67 |
| D | 363 | 63.09 | 0 | 0 | 0 | 0 | 0 | 0 | 0 | 58.88 | 87.1 |
| E | 385 | 50.44 | 0 | 0 | 0 | 0 | 0 | 0 | 0 | 100 | 100 |
| R | 388 | 0 | 0 | 0 | 0 | 50.44 | 0 | 0 | 0 | 97.38 | 100 |
| E | 391 | 0 | 0 | 0 | 0 | 0 | 1.99 | 0 | 0 | 92.9 | 93.83 |
| R | 394 | 0 | 0 | 1.99 | 0 | 0 | 0 | 0 | 0 | 91.59 | 100 |
| E | 402 | 0 | 0 | 80.48 | 0 | 0 | 0 | 0 | 0 | 100 | 100 |
| R | 404 | 0 | 0 | 0 | 0 | 0 | 80.48 | 0 | 0 | 96.45 | 96.45 |
| R | 415 | 0 | 0 | 0 | 0 | 61.68 | 0 | 0 | 0 | 16.82 | 90.09 |
| K | 418 | 0 | 0 | 0 | 0 | 1.41 | 0 | 0 | 0 | 51.59 | 86.92 |
| E | 436 | 0 | 20.02 | 0 | 0 | 0 | 0 | 0 | 0 | 28.22 | 49.35 |
| E | 440 | 0 | 78.3 | 0 | 0 | 0 | 0 | 0 | 0 | 30.65 | 56.26 |
| E | 444 | 0 | 1.65 | 0 | 0 | 0 | 0 | 0 | 0 | 38.13 | 69.16 |
| R | 451 | 0 | 14.6 | 0 | 0 | 0 | 0 | 0 | 0 | 36.82 | 61.5 |
| E | 458 | 0 | 47.74 | 0 | 0 | 0 | 0 | 0 | 0 | 85.23 | 93.27 |
| R | 468 | 0 | 0 | 0 | 0 | 0 | 0 | 14.23 | 0 | 49.91 | 70.47 |

| | | | | | | | | | | | |
|---|---|---|---|---|---|---|---|---|---|---|---|
| D | 471 | 0 | 0 | 0 | 0 | 0 | 0 | 0 | 26.04 | 15.33 | 43.93 |
| D | 481 | 0 | 0 | 0 | 95.86 | 0 | 0 | 0 | 0 | 87.85 | 93.27 |
| R | 482 | 0 | 0 | 0 | 0 | 0 | 0 | 0 | 164.33 | 17.57 | 32.71 |
| E | 503 | 0 | 0 | 4.17 | 0 | 0 | 0 | 0 | 0 | 52.34 | 77.94 |
| R | 507 | 0 | 0 | 1.82 | 0 | 0 | 0 | 0 | 0 | 16.67 | 43.45 |
| E | 510 | 0 | 0 | 111.61 | 0 | 0 | 0 | 0 | 0 | 17.98 | 40.82 |
| R | 516 | 0 | 0 | 0 | 0 | 0 | 49.29 | 0 | 0 | 45.13 | 73.6 |

Table S3: The frequency of the residues participating in inter-dimeric hydrogen bond formation between particular monomers and the residue conservation rate based on identity and similarity (CDEHKNQRST residues).

| Res | ID | Frequency, % | | | | | | | | Conservation, % | |
|---|---|---|---|---|---|---|---|---|---|---|---|
| | | C->E | C->F | D->E | D->F | E->C | E->D | F->C | F->D | Identity | Similarity |
| R | 265 | 0.03 | 0 | 0 | 0 | 13.69 | 0 | 0 | 0 | 10.47 | 82.62 |
| D | 269 | 13.69 | 0 | 0 | 0 | 0 | 0 | 0 | 0 | 11.21 | 80.56 |
| R | 271 | 0 | 0 | 0 | 0 | 0 | 7.6 | 0 | 0 | 72.90 | 95.33 |
| N | 272 | 0 | 0 | 0 | 0 | 0 | 10.32 | 0 | 0 | 9.91 | 88.04 |
| N | 275 | 0 | 0 | 0 | 0 | 0 | 2.56 | 0 | 0 | 5.23 | 79.63 |
| Q | 297 | 0 | 0 | 0 | 0 | 0 | 0.03 | 0 | 0 | 46.36 | 96.45 |
| E | 303 | 0 | 0 | 0.03 | 60.93 | 0 | 0 | 0 | 0 | 62.99 | 83.36 |
| Q | 311 | 0 | 0 | 0 | 2.49 | 0 | 0 | 0 | 0 | 71.78 | 99.44 |
| T | 315 | 0 | 0 | 0 | 0.13 | 0 | 0 | 0 | 0 | 86.92 | 93.83 |
| K | 317 | 0 | 0 | 0 | 0 | 0 | 0 | 0 | 1.08 | 66.36 | 95.14 |
| Q | 318 | 0 | 0 | 0 | 5.46 | 0 | 0 | 0 | 0 | 60.75 | 91.03 |
| E | 321 | 0 | 0 | 0 | 5.25 | 0 | 0 | 0 | 1.91 | 31.21 | 89.16 |
| N | 322 | 0 | 0 | 0 | 1.68 | 0 | 0 | 0 | 0 | 76.07 | 92.90 |
| R | 324 | 0 | 0 | 0 | 1.88 | 0 | 0 | 0 | 10.13 | 44.11 | 90.47 |
| Q | 325 | 0 | 0 | 0 | 8.33 | 0 | 0 | 0 | 0.13 | 54.95 | 92.34 |
| H | 328 | 0 | 0 | 0 | 0.07 | 0 | 0 | 0.54 | 0.33 | 2.80 | 90.65 |
| S | 332 | 0 | 0 | 0 | 0.03 | 0 | 0 | 0.84 | 0 | 21.87 | 91.03 |
| E | 335 | 0 | 0 | 0 | 0 | 0 | 0 | 18.88 | 0 | 19.44 | 80.56 |
| R | 339 | 0 | 0 | 0 | 0 | 0 | 0 | 8.98 | 0 | 20.93 | 93.83 |
| N | 346 | 0 | 0 | 0 | 0 | 0 | 0 | 0.03 | 0 | 20.75 | 83.18 |
| Q | 359 | 3.56 | 0 | 0 | 0 | 0 | 0 | 0 | 0 | 21.31 | 84.30 |
| D | 363 | 35.19 | 0 | 0 | 0 | 0 | 0 | 0 | 0 | 58.88 | 94.21 |
| S | 366 | 0.24 | 0 | 0 | 0 | 0 | 0 | 0 | 0 | 45.23 | 70.65 |
| Q | 374 | 0.3 | 0 | 0 | 0 | 0 | 0 | 0 | 0 | 100.00 | 100.00 |
| N | 376 | 0 | 0 | 0 | 0 | 0.3 | 0.14 | 0 | 0 | 98.50 | 100.00 |
| N | 381 | 0.16 | 0 | 0 | 0 | 0.16 | 0 | 0 | 0 | 100.00 | 100.00 |
| E | 385 | 0 | 0 | 0 | 0 | 0 | 0 | 0 | 0 | 100.00 | 100.00 |

| | | | | | | | | | | | |
|---|---|---|---|---|---|---|---|---|---|---|---|
| R | 388 | 0 | 0 | 0 | 0 | 0 | 12.25 | 0 | 0 | 97.38 | 100.00 |
| Q | 392 | 0 | 0 | 12.25 | 0 | 0 | 0 | 0 | 0 | 78.69 | 95.14 |
| E | 402 | 0 | 0 | 0.98 | 0 | 0 | 0 | 0 | 0 | 100.00 | 100.00 |
| R | 404 | 0 | 0 | 0 | 0 | 0 | 0.84 | 0 | 0 | 96.45 | 96.45 |
| Q | 408 | 0 | 0 | 0 | 0 | 0 | 1.14 | 0 | 0 | 79.81 | 96.82 |
| R | 409 | 0 | 0 | 1.17 | 0 | 0 | 0 | 0 | 0 | 90.28 | 100.00 |
| Q | 412 | 0 | 0 | 0 | 0 | 0 | 0.03 | 0 | 0 | 46.17 | 91.96 |
| R | 415 | 0 | 0 | 0 | 0 | 17.23 | 0 | 0 | 0 | 16.82 | 97.57 |
| S | 419 | 0 | 0 | 0 | 0 | 18.2 | 0 | 0 | 0 | 11.03 | 54.77 |
| E | 422 | 0 | 0 | 0 | 0 | 0.47 | 0 | 0 | 0 | 32.90 | 77.76 |
| D | 423 | 0 | 0 | 0 | 0 | 3.09 | 0 | 0 | 0 | 26.92 | 89.72 |
| E | 436 | 0.03 | 0.03 | 0 | 0 | 0 | 0 | 0 | 0 | 28.22 | 79.81 |
| S | 437 | 0 | 0 | 0 | 0 | 0.03 | 0 | 0 | 0 | 16.82 | 87.48 |
| E | 440 | 0 | 8.95 | 0 | 0 | 0 | 0 | 0 | 0 | 30.65 | 86.54 |
| S | 447 | 0 | 18.88 | 0 | 0 | 0 | 0 | 0 | 0 | 18.50 | 81.31 |
| R | 451 | 0 | 1.41 | 0 | 0 | 0 | 0 | 0 | 0 | 36.82 | 95.14 |
| D | 464 | 0 | 0 | 0 | 0 | 0 | 0 | 0 | 0.34 | 43.36 | 90.28 |
| S | 467 | 0 | 0 | 0 | 0 | 0 | 0 | 0 | 4.68 | 67.10 | 88.41 |
| R | 468 | 0 | 0 | 0 | 0 | 0 | 0 | 0 | 2.48 | 49.91 | 89.53 |
| D | 471 | 0 | 0 | 0 | 0 | 0 | 0 | 0 | 1.82 | 15.33 | 83.93 |
| Q | 472 | 0 | 0 | 0 | 0 | 0 | 0 | 0 | 2.42 | 88.97 | 98.32 |
| R | 482 | 0 | 0 | 0 | 0.07 | 0 | 0 | 0 | 53.5 | 17.57 | 82.06 |
| Q | 485 | 0 | 0 | 0 | 1.44 | 0 | 0 | 0 | 0.27 | 93.27 | 99.07 |
| Q | 486 | 0 | 0 | 0 | 0 | 0 | 0 | 0 | 8.67 | 92.34 | 98.32 |
| E | 503 | 0 | 0 | 10.42 | 0 | 0 | 0 | 0 | 0 | 52.34 | 90.65 |
| Q | 504 | 0 | 0 | 0.03 | 0 | 0 | 0 | 0 | 0 | 90.26 | 99.44 |
| S | 506 | 0 | 0 | 0.2 | 0 | 0 | 0 | 0 | 0 | 17.04 | 69.85 |
| R | 507 | 0 | 0 | 10 | 0 | 0 | 0 | 0 | 0 | 16.67 | 73.22 |
| E | 510 | 0 | 0 | 6.12 | 0 | 0 | 0 | 0 | 0 | 17.98 | 86.89 |
| R | 516 | 0 | 0 | 0 | 0 | 0 | 6.29 | 0 | 0 | 45.13 | 93.07 |
| Q | 519 | 0 | 0 | 0 | 0 | 0.03 | 0 | 0 | 0 | 8.05 | 53.56 |
